# Low-nanogram Fourier Transform Isotopic Ratio Mass Spectrometry of Proteins

**DOI:** 10.64898/2026.01.30.702838

**Authors:** Hassan Gharibi, Ana C. Jorge, Xuepei Zhang, Roman A. Zubarev

**Affiliations:** Division of Physiological Chemistry I, Department of Medical Biochemistry and Biophysics, Karolinska Institutet, SE-17 177 Stockholm, Sweden; Single Cell Proteomics Core Facility, Karolinska Institutet, SE-17 177 Stockholm, Sweden; FT-ICR and Structural Mass Spectrometry Laboratory, Faculdade de Ciencias, Universidade de Lisboa, 1749-016, Libsoa, Portugal; Biosystems and Integrative Sciences Institute, Faculdade de Ciencias, Universidade de Libsoa, 1749-016, Libsoa, Portugal; Chemical Proteomics Unit, Science for Life Laboratory (SciLifeLab), Stockholm, Sweden; Chemical Proteomics, Swedish National Infrastructure for Biological Mass Spectrometry (BioMS), Stockholm, Sweden; SciLIfeLab, SE-17 177 Stockholm, Sweden

## Abstract

Stable carbon and nitrogen isotope ratios are widely used in the life sciences to investigate diet, trophic interactions, and metabolic fluxes, but conventional isotope ratio mass spectrometry requires milligram-scale samples, limiting its applicability to small or rare biological specimens. Fourier Transform Isotopic Ratio Mass Spectrometry (FT IsoR MS) enables amino acid–resolved isotope analysis in a proteomics-compatible workflow and has previously been demonstrated at the microgram scale. Here, we assess the lower sample limit of FT IsoR MS by integrating it with single-cell proteomics–style sample preparation. Using human HeLa cells cultured in ^13^C-glucose–enriched and control media, we show that reliable relative δ^13^C measurements can be obtained from as few as 50 cells, corresponding to <10 ng of total protein, with a precision of approximately ±9‰. The observed amino acid–specific labeling patterns are metabolically coherent and consistent with bulk measurements, while smaller cell numbers (≤10 cells) do not yield statistically robust results. These findings establish the practical sensitivity threshold of FT IsoR MS at the low-nanogram level and demonstrate its suitability for isotope-resolved analyses of small cell populations, micro-organoids, and other low-input biological samples, thereby extending stable isotope analysis toward single-cell–scale applications.

## INTRODUCTION

Stable isotope ratios of biological elements hydrogen, carbon and nitrogen have long been used as biochemical tracers to infer dietary patterns and ecological interactions. There are many reasons in research to seek objective information on diet, including studies on animal ecology^1^ and archaeology^2^. Climate research also often relies on historical records inferred from animal diets^3^. In human health, there is a pressing need for more objective dietary information, as strong empirical diet-disease associations observed in controlled laboratory settings may be compromised when dietary assessments rely on self-reporting, which is not always reliable^4^. In many cases the object of isotopic analysis is proteins, including collagen extracted from bones^5^. Traditional isotope ratio mass spectrometry (IR MS) consumes at least one milligram of sample per isotope measured, which for rare or precious samples is a limiting factor. This limitation is based on the need to convert proteins to gas by combustion or pyrolysis, with subsequent losses in the sample preparation device. In contrast, novel Fourier Transform Isotope Ratio Mass Spectrometry (FT IsoR MS) requires only digestion of protein sample by trypsin or any other protease. The experimental set-up is compatible with the label-free proteomics LC-MS/MS experiment and microgram sensitivity has been demonstrated for amino-acid resolved analysis^6,7^. Yet even that sensitivity is insufficient for analysis of “micro-organoids” or spheroids that bridge the gap between 2D cell cultures and complex organoids or in vivo model: often they contain only 1,000–5,000 cells each^8^. With each cell providing ca. 100-150 pg of protein, the total amount can be as low as 100 ng. Therefore, higher sensitivity is required for FT IsoR MS analysis. The progress of single cell proteomics (SCP) with reliable detection of ≥5000 proteins in label-free analysis^9^ inspired us to investigate the minimal sample consumption in FT IsoR MS analysis using the SCP-style sample preparation. We thus investigated the ability of FT IsoR MS to correctly assess the isotope enrichment pattern when the sample consisted on N individual human HeLa cells (N = 1, 5, 10, and 50). The ^13^C enrichment pattern of several monitored amino acids relative to unlabelled samples was compared to that of the bulk FT IsoR MS analysis. To distinguish from the bulk, the samples with N≤50 cells are called here few-cell samples.

### EXPERIMENTAL SECTION

HeLa cells were originally obtained from ATCC®. Cells were cultured at 37 °C in Dulbecco’s Modified Eagle Medium (DMEM, Thermo Fisher Scientific, Cat. 11594446) supplemented with 10% heat-inactivated fetal bovine serum (FBS, Thermo Fisher Scientific, Cat. 11560636) and 1% penicillin-streptomycin (PenStrep, Gibco, Cat. 15140-122) in a humidified atmosphere containing 5% CO□. Isotope-labeled medium (10% ^13^C-glucose) and control medium were prepared by dissolving ^13^C-glucose (Silantes, ^13^C > 98.5%) or unlabelled glucose (Sigma, G7021) in glucose-free DMEM (Gibco, 11966025), along with 10% FBS and 1% penicillin-streptomycin as described above. Cells were washed twice with either ^13^C-glucose or control medium prior to seeding. After 7 weeks of culture, cells were harvested for further analysis.

Thereafter, cells were sorted by flow cytometry based on the forward and side scatter parameters in a Aria Fusion instrument (BD Biosciences). Either N=1, 5, 10, or 50 cells per well were collected in 96-well Lo-Bind plates (Eppendorf, Hamburg, Germany), with each well containing 5 μL of 100 mM triethylammonium bicarbonate (TEAB) pH 8.5 buffer.

For protein extraction, the plates underwent 5 freeze-thaw cycles, with freezing for 1 min in liquid nitrogen followed by immediate heating at 37 ºC for 1 min. For protein digestion, 1 µL of 25 ng/µL sequencing grade trypsin (Promega, Madison, WA) in 100 mM TEAB buffer was added to each well using MANTIS® liquid dispenser (Formulatrix, Bedford, MA), followed by overnight incubation at 37 ºC. The digested samples were dried in a SpeedVac (Concentrator Plus, Eppendorf) and stored at -20 ºC until further use.

For bulk samples, Hela cells were collected and washed with PBS twice. The cells were resuspended in 20 mM EPPS (Merck, E9502) buffer at pH 8.2 containing protease inhibitors (Thermo Fisher Scientific, Cat. 78439) and were subjected to three freeze/thaw cycles to lyse the cells. The protein concentration was measured using a rapid protein quantification assay kit (Thermo Fisher Scientific, Pierce 660LJnm) according to the manufacturer’s protocol. For each independent digest, approximately 50 µg of proteins was aliquoted and later subjected to digestion with trypsin at a 1:100 (protein/enzyme) ratio overnight. The resultant peptides were acidified by formic acid (FA) to a final concentration of 1% and desalted using C18 Sep-Pak cartridge. The eluates were collected, dried in a SpeedVac, and dissolved in a 2% ACN, 0.1% FA solution to a concentration of 0.5 µg/µL before injection into LC-MS/MS.

The FT IsoR MS analysis was performed as follows. Dried samples were incubated for 15 min at room temperature (RT) with 6 µL of buffer A (2% v/v acetonitrile, and 0.1% v/v formic acid in water). The peptide solution was transferred to the autosampler vials of the Nanoflow UltiMate 3000 UPLC coupled to an Orbitrap Fusion Eclipse Tribrid mass spectrometer equipped with a field asymmetric ion mobility spectrometry (FAIMS) Pro interface (all Thermo Fisher Scientific). Peptides were separated on a 50 cm Easy-Spray PepMap C-18 column (Thermo Fisher Scientific) at a temperature of 55 °C. The liquid phase consisted of buffer A and buffer B (2% v/v water, and 0.1% v/v formic acid in acetonitrile). The elution gradient ranged from 4% B to 15% B over 5 min with a flow of 0.3 µL/min, remained at 15% B for 1 min, with a flow of 0.1 µL/min, increased to 55% B for 71 min with a flow of 0.1 µL/min, increased to 95% with a flow of 0.1 µL/min, where it remained for 3 min at a flow of 0.3 µL/min. Finally, the buffer B was decreased to 4% at a 0.3 µL/min flow for 2 min, where it remained for 5 min before starting a new cycle. FAIMS operated at a compensation voltage of -43 V. Mass spectra were acquired in the positive ion mode in the Orbitrap analyzer using profile mode over an m/z range of 375-1500, with one microscan per mass spectrum. The nominal resolution was 120,000 and automatic gain control (AGC) was set to a standard target with a maximum injection time (IT) of 50 ms. The precursor ions for MS/MS were selected with a quadrupole isolation window of 1000 m/z and presumed charge state of 2+. Fragmentation was performed using higher-energy collisional dissociation (HCD) with a normalized collision energy of 50%. Ion detection was performed at a nominal resolution of 60,000 with m/z range between 50 and 200. The AGC target was set to 1000% with a 80 ms IT. The mass spectra were recorded in full profile mode, with 20 microscans per mass spectrum. All samples but bulk were analyzed in n≥3 independently obtained and processed biological (for bulk, 3 independent proteome digest) replicates. The replicate samples were analyzed in randomized order.

The acquired .raw files were converted to .mzML format using MSConvert (version 3.0.20168) from ProteoWizard. The .mzML files were then reformatted to a .csv table using the in-house PAIR-MS (https://github.com/hassanakthv/PAIR-MS) github package. The isotopic ratios were determined as explained in Gharibi et al^7,10^.

## RESULTS

In HeLa cell analysis, isotopic ratios for the following amino acids were monitored via immonium ions: leucine and isoleucine (L/I, indistinguishable in FT IsoR MS), proline (P), pyroglutamate (Pyr), and valine (V). Despite the fact that the HeLa cells were cultured for seven weeks in the media containing 10% ^13^C-glucose, bulk FT IsoR MS analysis showed that only two amino acids were enriched, P and Pyr, and to a modest degree (≈65‰ compare to isotopically normal control), while L/I and V showed no statistically significant difference (Figure 1).

**Figure 1.**
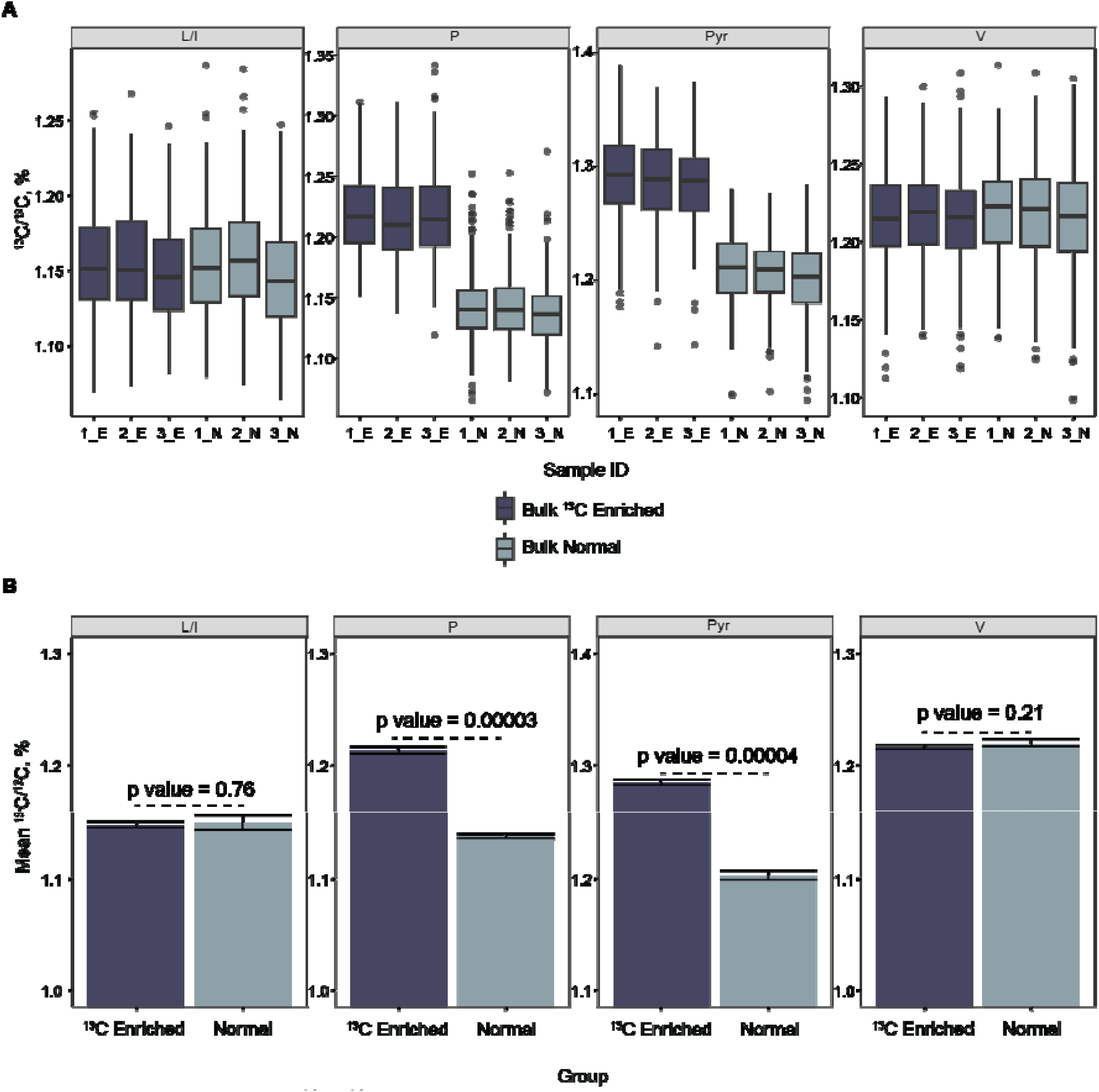
Overview of ^13^C/^12^C measurement in the HeLa bulk experiment. (A) Boxplots showing the ^13^C/^12^C distribution for L/I, P, Pyr, and V amino acids for each sample. (B) Mean ^13^C/^12^C ratios across three replicates for cells grown in normal and ^13^C-enriched media (error bars represent the standard deviation of three replicates).

For 50 cell analysis, a similar pattern of isotope enrichment as in bulk measurements was observed (Figure 2).

**Figure 2.**
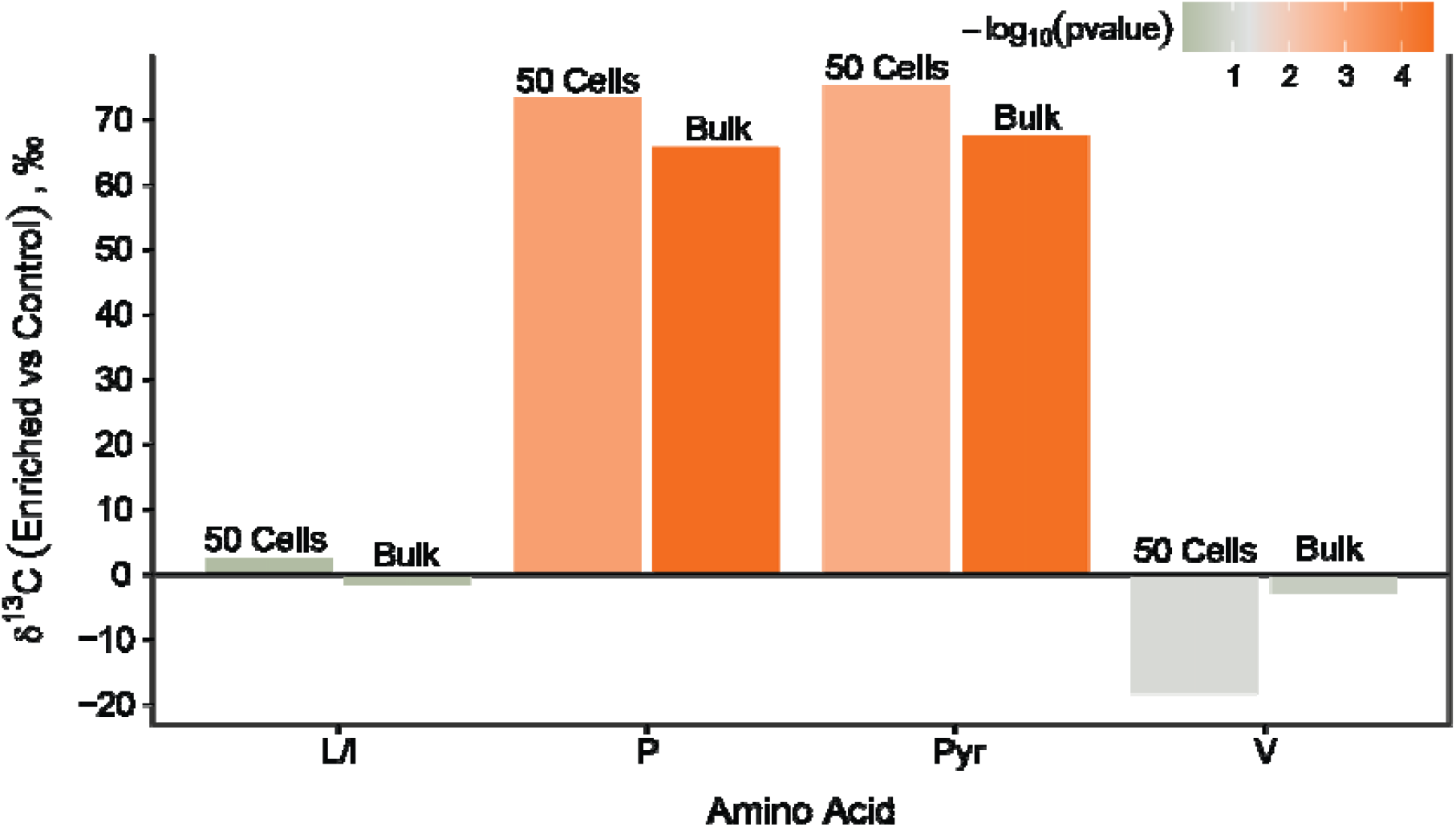
Bar plot showing the δ^13^C values (using Control as the reference) for abundant amino acids for both 50-cell (n = 5) and bulk (n=3) HeLa samples.

The observed δ^13^C difference between the enriched and normal groups reached high statistical significance, with p-value for P was 0.0007 for 50 cells, compared to p = 0.00003 obtained in the bulk analysis. For Pyr, the results were similar. The isotopic ratios in individual replicates were also consistent with those observed in bulk samples (Figure 3).

**Figure 3.**
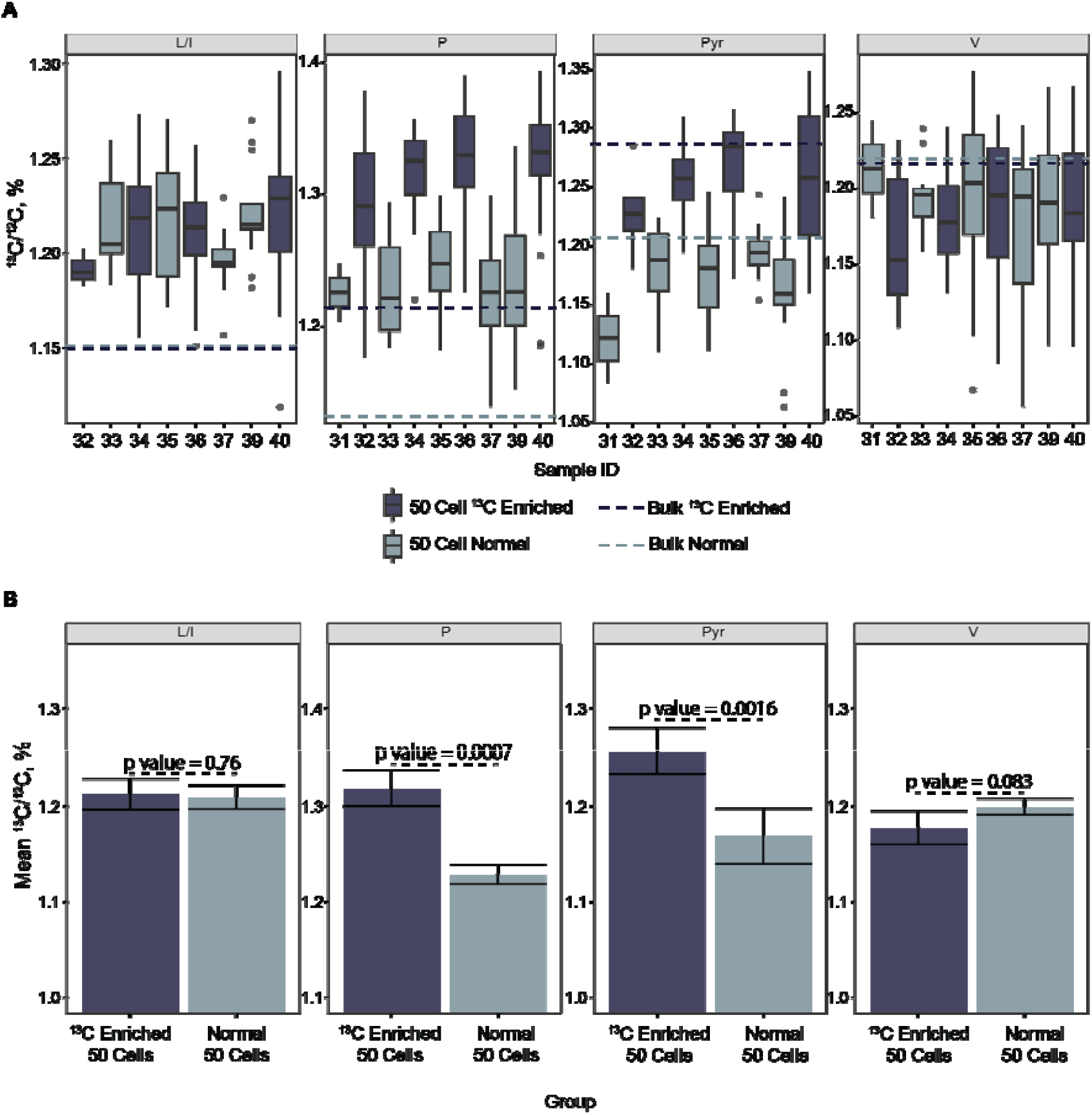
Overview of ^13^C/^12^C measurements in a 50-cell HeLa experiment. (A) Boxplots of the ^13^C/^12^C distribution for amino acids L/I, P, Pyr, and V in different replicate analyses. (B) Mean ^13^C/^12^C ratios across three replicates for cells grown in normal and ^13^C-enriched media. Error bars represent the standard deviation between the replicates.

For other tested cell numbers (1, 5, and 10 cells), the pattern and magnitudes of observed enrichments differed significantly from bulk measurements, and the differences between treatments haven’t reached statistical significance (Supplementary Figure 1).

## Discussion

Incorporation of ^13^C in pyroglutamate and proline from the isotopically labelled glucose was significant in comparison to other monitored amino acids. That enrichment pattern is metabolically coherent and expected. As labelled carbon enters central metabolism via glycolysis and the TCA cycle, ^13^C from glucose is efficiently incorporated into TCA intermediates, α-ketoglutarate becomes ^13^C-labeled, and transamination produces labelled glutamate. Pyroglutamic acid (5-oxoproline) is a cyclized derivative of glutamate (often via glutathione or spontaneous cyclization),^11^ so labelling here is also expected. The cyclization can either occur spontaneously or enzymatically by the action of glutaminyl cyclase^12,13,14^. Proline, a non-essential amino acid, is also mainly synthesized from glutamate. Besides, proline can be generated from arginine and ornithine, which also have glutamate as precursor^15,16^. Thus ^13^C-labeling of pyroglutamate and proline is justified by their precursor, ^13^C-glutamate, derived from ^13^C-glucose. As the supplemented medium in which cells grew also had non-labelled glutamine (584 mg/L) that could also be converted into glutamate by the cells, the degree of ^13^C enrichment was small.

On the other hand, as valine, leucine, and isoleucine are branched-chain essential amino acids (BCAAs) in humans and cannot be synthesized *de novo*, HeLa cells import them directly from the medium that contained them in unlabelled form. Thus, in HeLa proteins these BCAAs maintained natural abundance of heavy carbon regardless of how much ^13^C-labelled glucose was supplied.

In the 50-cell analysis, the ^13^C/^12^C ratios directly measured from the MS/MS spectra were somewhat shifted compared to those from the bulk (Figure 3): for instance, for L/I in unlabelled samples ^13^C/^12^C was on average 1.15% for bulk and 1.20% for 50 cells. This corresponds to the relative difference of 44‰. However, the corresponding difference in δ^13^C was much smaller, ≤9 ‰ (Figure 2). This result highlights the fact that, even though direct isotope ratios measurements (^13^C/^12^C) in FT IsoR MS produce reasonable results, relative measurements compared to a standard (δ^13^C) give more accurate data.

## CONCLUSIONS

Here, using the workflow from single cell proteomics we investigated the minimal sample requirements in FT isoR MS. We found that 50 human HeLa cells with a total protein content below 10 nanogram produce data consistent with bulk analysis within <10‰. The 60-70‰ enrichment degree found in labelled P and Pyr amino acid residues is comparable with the range of heavy carbon enrichment in nature, approximately from -60‰ to +10‰. Therefore, not only the degree of artificial labelling, but also large natural variations in heavy carbon content can be measured by FT IsoR MS biological samples at a low-nanogram level.

The ability to measure amino acid-resolved carbon isotope ratios from <10 ng of total protein establishes FT IsoR MS as a viable analytical approach for samples previously inaccessible to isotope analysis. This sensitivity places FT IsoR MS within the material regime of single-cell proteomics and enables isotope-resolved measurements in small cell populations, micro-organoids, spatially microdissected tissues, and rare or precious biological samples. In such contexts, relative δ^13^C measurements referenced to internal or external standards are expected to be more robust than absolute isotope ratios and are well suited for comparative studies across conditions, time points, or microenvironments. Coupling FT IsoR MS with established single-cell and low-input proteomics workflows opens opportunities to investigate metabolic labelling patterns, nutrient routing, and isotopic heterogeneity at previously inaccessible spatial and biological scales, thereby extending stable isotope analysis from bulk tissues to the level of small cellular assemblies.

## ASSOCIATED CONTENT

### Data Availability Statement

The supporting information and files are available free of charge upon request.

## Acknowledgements

This work was supported by the EU FET OPEN grant ARIADNE VIBE and Swedish Research Council grant 2021-05223.

---

**Supplementary Figure 1.**
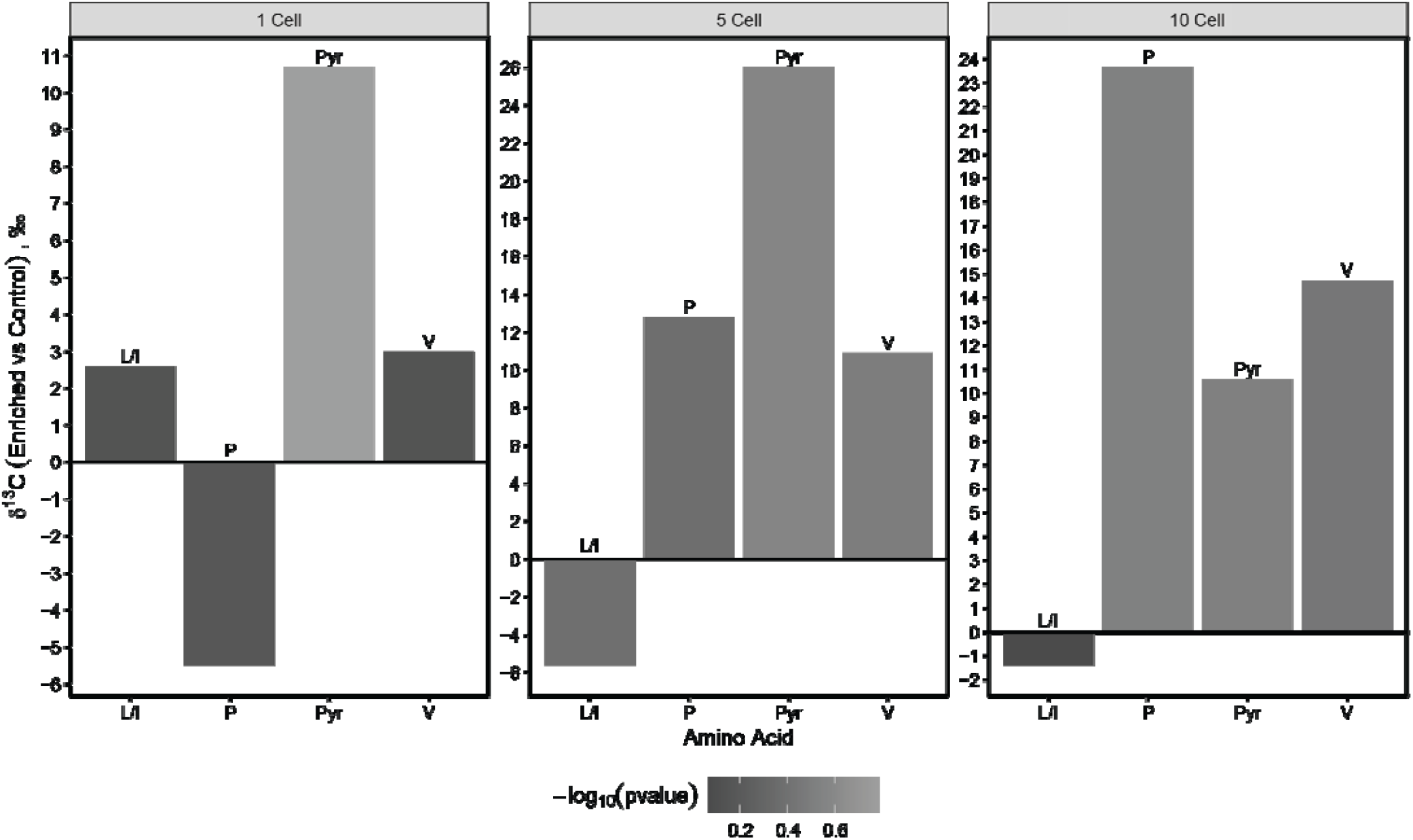
Bar plot showing the δ^13^C values (using Control as the reference) in 1, 5, and 10-cell HeLa experiments.

